# CRISPR-Cas9 screening of KSHV-transformed cells identifies XPO1 as a vulnerable target of cancer cells

**DOI:** 10.1101/601369

**Authors:** Marion Gruffaz, Hongfeng Yuan, Wen Meng, Hui Liu, Sangsu Bae, Jin-Soo Kim, Chun Lu, Yufei Huang, Shou-Jiang Gao

**Author notes:** Address correspondence to Shou-Jiang Gao.

## Abstract

The abnormal proliferation of cancer cells is driven by deregulated oncogenes or tumor suppressors, of which the cancer vulnerable genes are attractive therapeutic targets. Targeting mislocalization of oncogenes and tumor suppressors resulting from aberrant nuclear export is effective for inhibiting growth transformation of cancer cells. We performed a CRISPR-Cas9 screening in a unique model of matched primary and oncogenic KSHV-transformed cells, and identified genes that were pro-growth and growth-suppressive of both cells, of which exportin XPO1 was demonstrated to be critical for the survival of transformed cells. Using XPO1 inhibitor KPT-8602 and by siRNA knockdown, we confirmed the essential role of XPO1 in cell proliferation and growth transformation of KSHV-transformed cells, and cell lines of other cancers including gastric cancer and liver cancer. XPO1 inhibition induced cell cycle arrest through p53 activation but the mechanism of p53 activation differed among different types of cancer cells. p53 activation depended on the formation of PML nuclear bodies in gastric cancer and liver cancer cells. Mechanistically, XPO1 inhibition induced relocalization of autophagy adaptor protein p62 (SQSTM1), recruiting p53 for activation in PML nuclear bodies. Taken together, we have identified novel pro-growth and growth-suppressive genes of primary and cancer cells, and demonstrated XPO1 as a vulnerable target of cancer cells. XPO1 inhibition induces cell arrest through a novel PML-and p62-dependent mechanism of p53 activation in some types of cancer cells.

**Importance:** Using a model of oncogenic virus KSHV driven cellular transformation of primary cells, we have performed a genome-wide CRISPR-Cas9 screening to identify vulnerable genes of cancer cells. This screening is unique in that this virus-induced oncogenesis model does not depend on any cellular genetic alterations, and has matched primary and KSHV-transformed cells, which are not available for similar screenings in other types of cancer. We have identified genes that are both pro-growth and growth-suppressive in primary and transformed cells, some of which could represent novel proto-oncogenes and tumor suppressors. In particular, we have demonstrated exportin XPO1 as a critical factor for the survival of transformed cells. Using a XPO1 inhibitor KPT-8602 and by siRNA-mediated knockdown, we have confirmed the essential role of XPO1 in cell proliferation and growth transformation of KSHV-transformed cells, as well as gastric and liver cancer cells. XPO1 inhibition induces cell cycle arrest by activating p53 but the mechanism of p53 activation differed among different types of cancer cells. p53 activation is dependent on the formation of PML nuclear bodies in gastric and liver cancer cells. Mechanistically, XPO1 inhibition induces relocalization of autophagy adaptor protein p62 (SQSTM1), recruiting p53 for activation in PML nuclear bodies. These results illustrate XPO1 as a vulnerable target of cancer cells, and reveal a novel mechanism for blocking cancer cell proliferation by XPO1 inhibition as well as a novel PML-and p62-mediated mechanism of p53 activation in some types of cancer cells.

## Introduction

The malfunction of nuclear transport, which shuttles proteins between cytoplasm and nucleus, often leads to mislocalization of oncogenes and tumor suppressor proteins in cancer cells(1). Indeed, numerous proteins involved in cancer including p53, APC, Retinoblastoma (Rb), NFAT and β-catenin are abnormally localized in cancer cells. The consequence of these dysfunctions is either over-activation of oncogenes, or inactivation of tumor suppressor proteins, resulting in uncontrolled cell proliferation and growth transformation(2).

Unlike passive diffusion of metabolites, shuttling of proteins between nucleus and cytoplasm is an active, highly regulated receptor-mediated process. The nuclear pore complex (NPC) is a supramolecular structure composed of more than 30 nucleoporins interacting with importins and exportins within the NPC channel. The directionality of the transport depends on the distribution of GTP-or GDP-bound GTPase, the Ran protein, in both cytoplasmic and nucleic compartments(3). Indeed, while importins bind to their cargo proteins in cytoplasm and release them in nucleus upon binding of RanGTP, exportins bind to cargos in nucleus only in the presence of RanGTP and release them in cytoplasm upon Ran-driven GTP hydrolysis(4).

The exportin family consists of 7 members including XPO1 (CRM1), XPO2 (CSE1L), XPO3 (XPOt), XPO4, XPO5, XPO6 and XPO7. While XPO1, XPO2, XPO4, XPO6 and XPO7 primarily mediate the export of cargo proteins, XPO3 and XPO5 are involved in the transport of tRNAs and precursor microRNAs (pre-miRNAs), respectively. XPO1, the major protein export receptor, is also associated with the nuclear export of mRNAs and rRNAs(5).

Mechanistically, XPO1 interacts with the nucleoporins NUP214 and NUP88 inside the NPC and exports proteins containing XPO1-specific nuclear export signal (NES), which are short leucine-rich sequences. Several tumor suppressor proteins and oncogenes display XPO1-specific NES(6). Consequently, XPO1 dysregulation indirectly regulates cellular functions including cell proliferation and cellular transformation, apoptosis, and chromosome segregation(2). Particularly, XPO1 is upregulated in ovarian carcinoma, glioma, osteosarcoma, pancreatic, cervical and gastric cancers, which induces abnormal accumulation of the tumor suppressor proteins Rb, APC, p53, p21 and p27 in the cytoplasm, leading to their losses of nuclear functions(2). On the other hand, inhibition of XPO1 by RNA interference or with inhibitors, such as Leptomycin B or selective inhibitors of nuclear export (SINE), prevents cellular transformation and tumorigenesis in numerous cancer models. Hence, XPO1 has become a promising target in cancer therapy(7).

Nuclear bodies (NBs) are membraneless structures within the nucleus involved in multiple pathways of genome maintenance. Among them, promyelocytic leukemia protein nuclear bodies (PML-NBs) are involved in DNA repair, DNA damage response, telomere homeostasis, and p53-associated apoptosis and cell cycle arrest(8). Of the various biological consequences of XPO1 inhibition, several groups observed the retention of p53, the most described tumor suppressor, in the nucleus, enhancing p53-mediated tumor suppressor activity(8). Indeed, activation of p53 induces cell cycle arrest and apoptosis in many types of cancer cells. Particularly, inside the nucleus, p53 colocalizes and interacts with PML-NBs, which serve as p53-coactivator(9). PML knockout impairs p53-dependent apoptosis, p53-mediated transcriptional activation, as well as induction of p53 target genes such as Bax and p21(9). Therefore, PML-NBs may play a significant role in apoptosis and cancer. Nevertheless, the underlying molecular mechanism mediating PML-dependent p53 activation is still unclear.

The development of new genome engineering technologies has enabled the identification of oncogenes and tumor suppressors in cancer cells in vitro and in vivo(10, 11). Particularly, the clustered regularly interspaced short palindromic repeats (CRISPR)-CRISPR-associated (Cas) proteins system, adapted to mammalian cells based on a mechanism of adaptive immunity of bacteria and archea, enhances the accessibility of genome manipulation by allowing the targeting of genes with specific RNA sequences(12). Briefly, CRISPR relies on Cas9 guided by sgRNAs (CRISPR RNAs) to induce loss of function (LOF) mutations via frameshifts in the coding region, leading to gene inactivation. CRISPR-Cas9 system has enabled different types of genetic modifications, such as gene disruption or transcriptional activation. Several types of biological screens based on the CRISPR-Cas9 system have already been carried out to identify viral restriction factors, oncogenes, and tumor suppressors, as well as develop T-cell immunotherapies.

In this study, by performing a genome-wide CRISPR-Cas9 screening of cells transformed by an oncogenic virus Kaposi’s sarcoma-associated herpesvirus (KSHV), we have identified cellular genes that are essential for cellular transformation(13). Briefly, a CRISPR pooled libraries containing sgRNAs that specifically target all known cellular genes were transduced into Cas9-expressing KSHV-transformed primary rat mesenchymal embryonic stem cells (KMM) and control primary rat mesenchymal stem cells (MM)(13). Genomic DNA from surviving MM and KMM cells at day 1, 4, 11 and 21 post-transduction were collected and sequenced, and the results were analyzed for the gain or loss of sgRNAs. We identified the exportin family members XPO1, XPO2, XPO3, XPO5 and XPO7 as the essential factors involved in the survival of KMM cells. We confirmed the essential role of XPO1 in cell proliferation and growth transformation using cell lines of other types of cancer, including AGS derived from gastric cancer and HUH7 derived from liver cancer. We showed that inhibition of XPO1 by siRNAs or with a XPO1 inhibitor KPT-8602 blocked cell proliferation and growth transformation by inducing p53-mediated cell cycle arrest in all types of cancer cells tested. However, we found that the mechanism mediating p53 activation differed among different types of cancer cells. In AGS and HUH7 cells, p53 activation was correlated with nuclear accumulation of autophagy adaptor protein p62 (SQSTM1), which was colocalized with p53 within the PML-NBs. Knockdown of p62 with siRNAs abrogated p53 activation induced by XPO1 inhibition. These results highlight an essential role of p62 in controlling cell proliferation and growth transformation by mediating activation of the p53 pathway in some types of cancer cells.

## Results

### Genome-wide CRISPR-Cas9 screening for essential genes of KSHV-transformed cells

KSHV is an oncogenic virus, which regulates numerous pro-growth and -survival pathways(14). We have previously shown that infection by KSHV alone is sufficient to efficiently infects and transforms MM cells without depending on cellular genetic alterations, and that KSHV-transformed KMM cells can efficiently induce tumors in nude mice with pathological features resembling Kaposi’s sarcoma (KS)(13). Because of the unique features of KSHV-induced cellular transformation and the available matched primary MM cells, KMM cells are ideal for identifying essential cellular genes that mediate growth transformation.

We first generated MM and KMM cells stably expressing Cas9 by lentiviral transduction following by positive selection with blasticidin for 1 week. Interestingly, we repeatedly observed weaker expression of Cas9 protein in MM than KMM cells in multiple experiments (Supplemental Fig. 1A). This was likely due to a higher rate of cell proliferation of KMM cells than MM cells, often leading to higher expression levels of genes(13, 15). Despite the differential levels of Cas9 expression, we observed efficient inhibition of the endogenous SIRT1 expression after lentiviral transduction of sgRNAs targeting SIRT1 in both Cas9-expressing MM and KMM cells (Supplemental Fig. 1B). Importantly, Cas9 expression did not affect the efficiency of colony formation of KMM cells in softagar (Supplemental Fig. 1C). As expected, Cas9-expressing MM cells did not form any colony in softagar(13). These results indicate that the Cas9-expressing MM and KMM cells can be used for identifying essential cellular genes that mediate growth transformation.

We then generated a library of sgRNAs targeting 19,840 genes of the rat genome, each with 3 independent sgRNAs. Cas9-expressing MM and KMM cells were transduced with the lentiviral sgRNA library and selected for 3 days with puromycin. The cultures were then switched to normal medium without any selection and cell samples were collected at day 1, 4, 11 and 21, and subjected to DNA sequencing to determine the gain and loss sgRNAs over time (Fig. 1A). Cumulative frequencies of sgRNAs were analyzed at day 1, 4, 11, and 21 post-transduction in MM and KMM cells (Fig. 1B). Compared to day 1, we observed a progressive shift to the left in the curves at day 4, 11, and 21 post-transduction, indicating depletion of a subset of sgRNAs. By assessing the global gene vulnerability in at day 4, 11, and 21 post-transduction, we observed a lower abundance of essential genes, displaying as negative CRISPR scores, in KMM than in MM cells, suggesting that KMM cells had higher survival rates than MM cells (Fig. 1B), which was also shown by the CRISPR scores of individual genes (Fig. 1C). Interestingly, the correlation between CRISPR score and gene expression at RNA level(16) was neither observed in MM nor in KMM cells (Supplemental Fig. 2). Finally, GSEA analysis at day 21 vs day 1 post-transduction revealed enrichments of gene signatures in housekeeping pathways including DNA replication, transcription and translation, which were significantly higher in KMM than in MM cells, confirming that MM cells were more susceptible to gene disruption than KMM cells (Fig. 1D).

**FIG 1.**
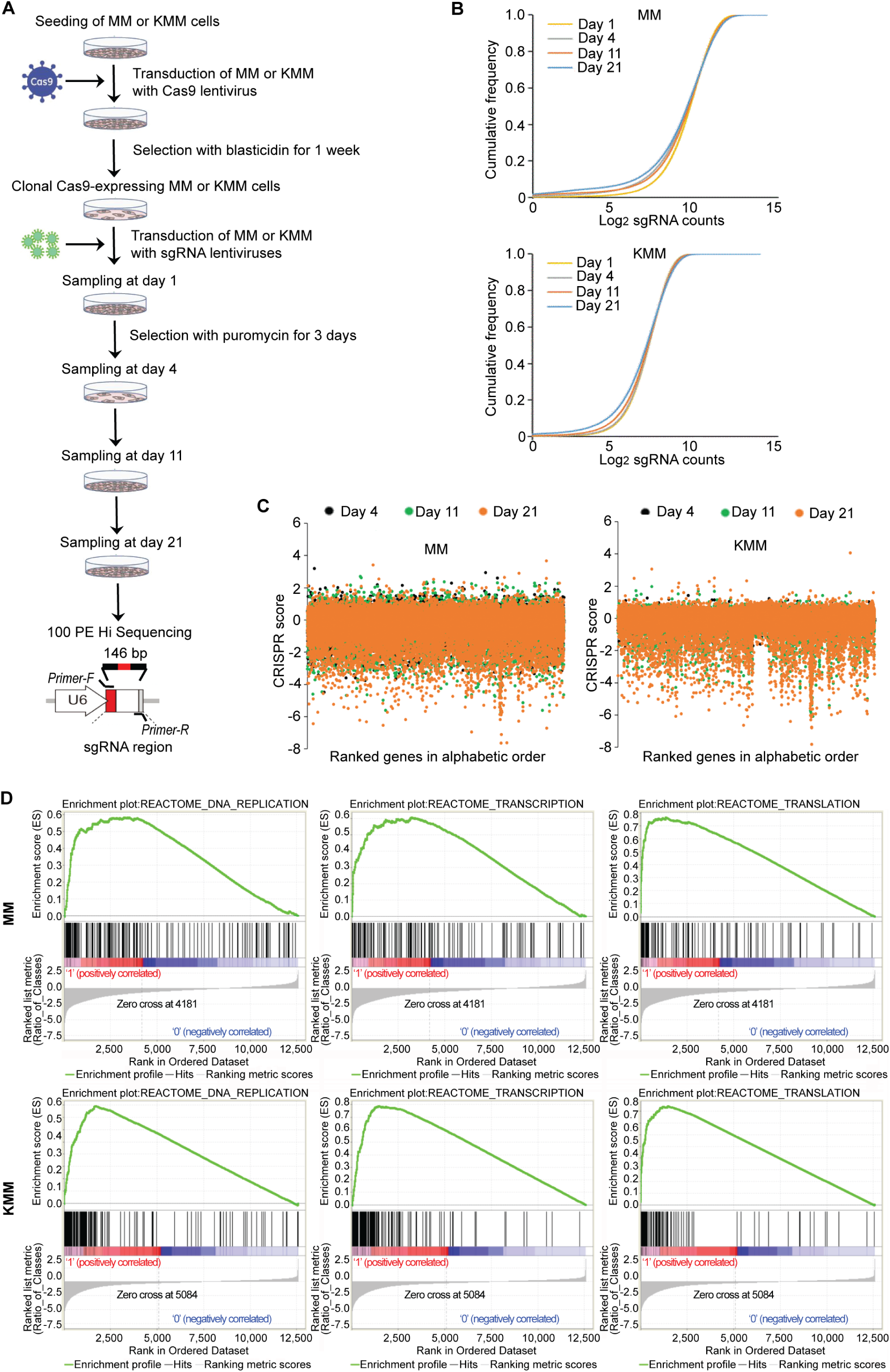
Genome-wide CRISPR-Cas9 screening for essential genes of KSHV-transformed cells. (A) Experimental design of CRISPR-Cas9 high throughput screening in KSHV-transformed cells (KMM), and matched primary MM cells. (B) Analysis of the cumulative frequency of sgRNAs at day 1, 4, 11, and 21 post-transduction in MM and KMM cells. (C) Analysis of gene essentiality at day 4, 11 and 21 post-transduction using CRISPR score in MM and KMM cells. Genes are ranked by alphabetical order. (D) GSEA analysis of enriched pathways at day 21 vs day 1 post-transduction in MM and KMM cells.

### Identification of essential genes and pathways of KSHV-transformed cells by CRISPR-Cas9 screening

Since essential genes are likely to encode key regulators of cellular processes involved in the survival of cancer cells, several studies investigated the overlapping of essential genes across different cancer cell lines by CRISPR-Cas9 screening(17). Therefore, we correlated results of MM and KMM cells with those of previous studies in chronic myeloid leukemia cell lines (KBM7 and K562) and Burkitt’s lymphoma cell lines (Raji and Jiyoye). We found a high degree of overlap in gene essentiality between KSHV-transformed cells and other types of cancer cells (R^2^≥0.29) (Fig. 2A). On the other hand, the correlation of essential genes remained low between the primary MM cells and other types of cancer cells (R^2^≤0.09).

**FIG 2.**
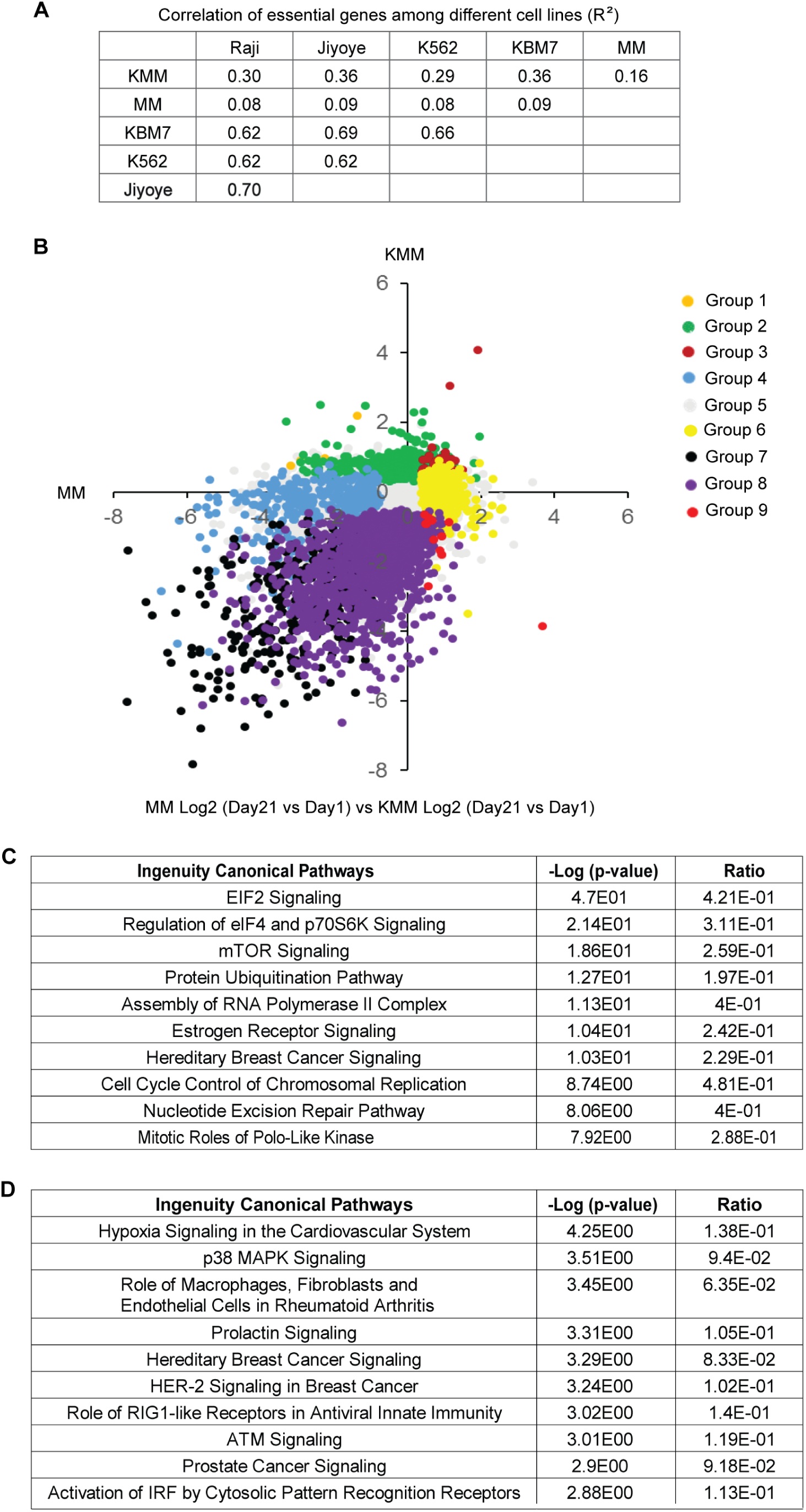
Identification of essential genes and pathways of KSHV-transformed cells. (A) Correlation of essential genes among MM, KMM, Raji, Jiyoye, K562 and KBM7 cell lines (R). (B) Classification of the 19,840 genes into 9 different groups based on the ratio of CRISPR score at day 21 over day 1 and statistical significance in MM versus KMM cells. Group 1 includes genes with KMM Log2 CRISPR score >0, p-value <0.05, and MM Log2 CRISPR score <0, p-value <0.05; Group 2 includes genes with KMM Log2 CRISPR score >0, p-value <0.05, and MM Log2 CRISPR score of p-value ≥0.05; Group 3 includes genes with both MM and KMM Log2 CRISPR scores >0, p-value <0.05; Group 4 includes genes with KMM Log2 CRISPR score of p-value ≥0.05, and MM Log2 CRISPR score <0, p-value <0.05; Group 5 includes genes with both MM and KMM Log2 CRISPR scores of p-value ≥0.05; Group 6 includes genes with KMM Log2 CRISPR score of p-value ≥0.05, and MM Log2 CRISPR score >0, p-value <0.05; Group 7 includes genes with both MM and KMM Log2 CRISPR scores <0, p-value <0.05; Group 8 includes genes with KMM Log2 CRISPR score <0, p-value <0.05, and MM Log2 CRISPR score of p-value ≥0.05; Group 9 includes genes with KMM Log2 CRISPR score <0, p-value <0.05, and MM Log2 CRISPR score >0, p-value <0.05. (C) IPA analysis of top 10 depleted pathways in KMM over MM cells at day 21 versus day 1 post-transduction. (D) IPA analysis of top 10 enriched pathways in KMM over MM cells at day 21 versus day 1 post-transduction.

We next classified the 19,840 genes based on CRISPR scores. For each type of cells, we first selected genes with CRISPR scores that were statistically different at day 21 over day 1, then between KMM and MM cells, and obtained 9 groups of genes (Fig. 2B and Table S1). Group 1 consisted of 11 genes that had significant increase in CRISPR score for KMM cells but significant decrease in CRISPR score for MM cells; Group 2 consisted of 680 genes that had significant increase in CRISPR score for KMM cells but no significant change in CRISPR score for MM cells; Group 3 consisted of 51 genes that had significant increases in CRISPR score for both KMM and MM cells; Group 4 consisted of 402 genes that had no significant change in CRISPR score for KMM cells but had significant decrease in CRISPR score for MM cells; Group 5 consisted of 16,558 genes that had no significant change in CRISPR score for both KMM and MM cells; Group 6 consisted of 552 genes that had no significant change in CRISPR score for KMM cells but had significant increase in CRISPR score for MM cells; Group 7 consisted of 314 genes that had significant decreases in CRISPR score for both KMM and MM cells; Group 8 consisted of 1,259 genes that had significant decrease in CRISPR score for KMM cells but had no significant change in CRISPR score for MM cells; and Group 9 consisted of 13 genes that had significant decrease in CRISPR score for KMM cells but had significant increase in CRISPR score for MM cells.

Of these 9 groups, genes in Group 8 were likely essential or pro-oncogenic for KMM cells but not for MM cells, of which, 18 genes had CRISPR score ratios of <-5 (−32 fold) at day 21 over day 1 for KMM cells, including Naa38, Rpl9_like, Rpl23a, Spcs3, Hspa14_like, Nfyb, Eif2b2, Mrpl55, Pold1, Nup43, Lin52, Csnk1a1, Aldoa, Rpl6, Ddx6, Wdr74, Rars and Cnot1. Genes in Group 2 were likely growth-suppressive for KMM cells but not MM cells. On the other hand, genes in Group 4 were likely essential or pro-growth for MM cells but not KMM cells while genes in Group 6 were likely growth-suppressive for MM cells but not KMM cells, suggesting that KSHV might target these two sets of genes to allow the transformed cells to overcome the essential or growth-suppressive functions of these genes, respectively.

We determined the molecular pathways that were significantly depleted and enriched in KMM over MM cells (day 21 vs day 1) by Ingenuity Pathway Analysis (IPA) (Supplemental Table 2 and 3). We found that the EIF2 pathway, the regulation of eIF4 and p70S6K, and the mTOR pathway were the top 3 depleted pathways in KMM cells, compared to MM cells, highlighting their roles in the survival of the transformed cells (Fig. 2C). In fact, these pathways are highly related to one another, and the mTOR pathway is the most effective target for the treatment of KS(18), hence further validating the relevance of this model for KS. On the other hand, we found the hypoxia pathway and the p38 MAPK pathway as the most enriched pathways in KMM cells compared to MM cells, suggesting their essential roles in maintaining the homeostasis of the KSHV-transformed cells (Fig. 2D).

### Identification of XPO1 as a critical factor involved in cell proliferation and growth transformation of cell lines derived from multiple types of cancer

Since oncogenes and tumor suppressor proteins are often mislocalized as a result of dysregulation of nuclear transport(1), we focused on the exportin family members. The counts of sgRNAs of XPO1, XPO2, XPO3 and XPO5 significantly decreased at day 21 compared to day 1 in KMM cells indicating the essential functions of these exportins, whereas XPO7 sgRNA counts significantly increased in KMM cells indicating its putative anti-survival effect in the transformed cells (Supplemental Fig. 3). In addition, MM cells were sensitive to XPO1, XPO2 and XPO5 knockout (Supplemental Fig. 3). Numerous inhibitors have been developed for the main exportin member, XPO1. Some of these inhibitors are currently in clinical trials for different types of cancer(19-21). Hence, we decided to focus on this exportin.

Since XPO1 is upregulated in numerous types of cancer(22), we first examined XPO1 expression in KMM cells compared to MM cells. We observed 3-fold higher expression level of XPO1 in KMM than MM cells at mRNA level (Fig. 3A), which was confirmed at protein level (Fig. 3B). Because KMM cells are latently infected by KSHV, expressing viral latent genes LANA, vFLIP, vCyclin and the miRNA cluster(13, 23), we examined the roles of these genes in XPO1 expression. Deletion of either vCyclin or the miRNA cluster but not vFLIP abolished XPO1 upregulation (Fig. 3A and 3B). Because LANA is essential for KSHV episome persistence, we were not able to obtain cells latently infected by the LANA mutant(14). However, overexpression of LANA in MM cells did not alter XPO1 expression (results not shown). By complementing cells infected by the miRNA cluster deletion mutant (ΔmiR) with individual miRNAs, we found that pre-miR-K3 mediated XPO1 upregulation (Fig. 3C). Hence, both vCyclin, a homologue of the cellular Cyclin D2(15), and pre-miR-K3, which activates the Akt pathway(24), not only promote cell cycle progression and facilitate G1/S phase transition, respectively (15, 23), but also cause XPO1 upregulation.

**FIG 3.**
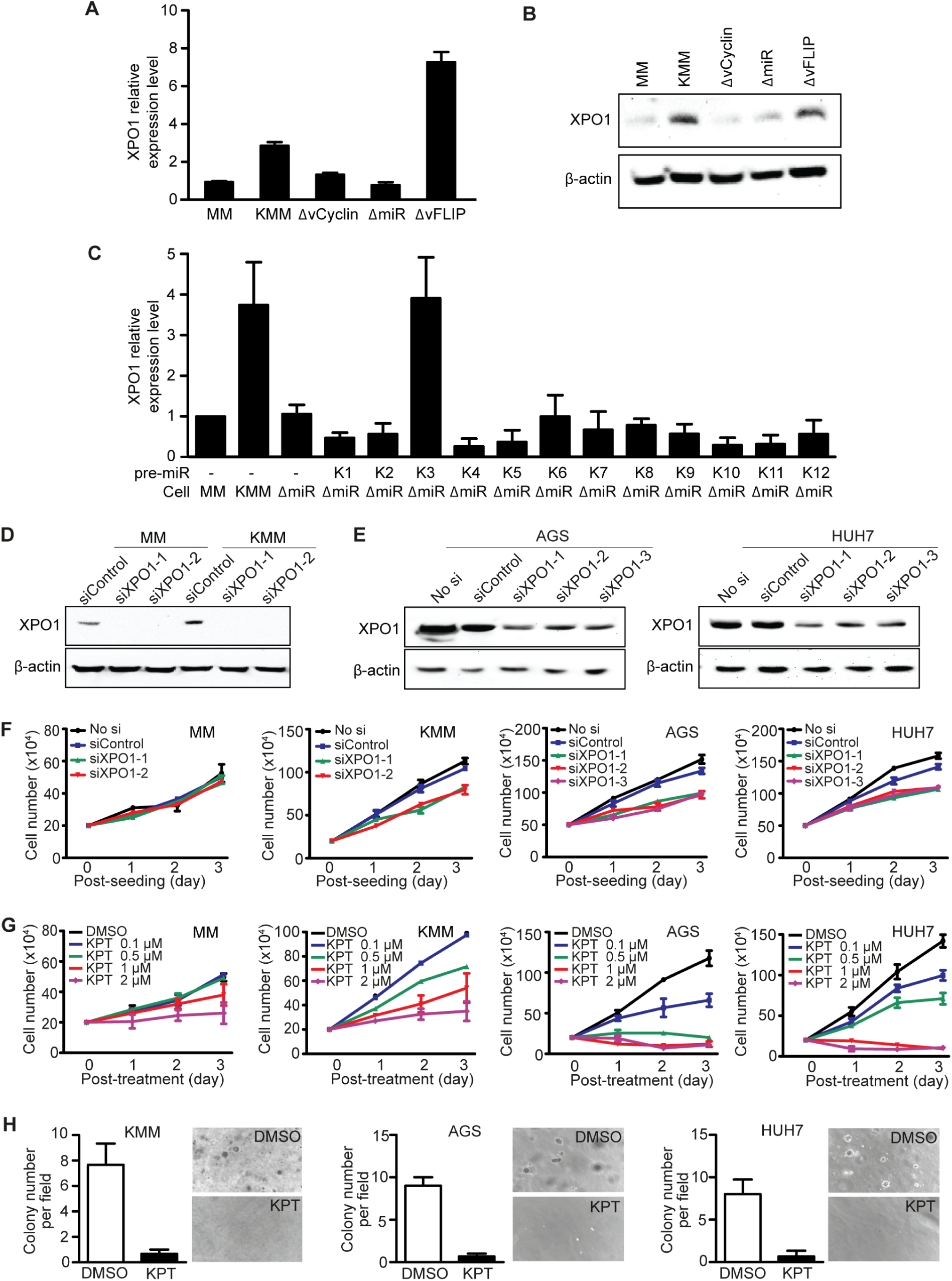
Identification of XPO1 as a vulnerable gene of KSHV-transformed KMM cells, and cell lines of gastric cancer AGS and liver cancer HUH7. (A-B) XPO1 expression in MM cells, and MM cells infected by KSHV (KMM), and mutant viruses with a deletion of vFLIP (ΔvFLIP), vCyclin (ΔvCyclin), or a cluster of 10 pre-miRNAs (ΔmiR) analyzed by qRT-PCR (A) and Western-blotting (B). (C) Expression of XPO1 analyzed by qRT-PCR in MM, KMM, ΔmiR cells, or ΔmiR cells complemented with individual KSHV pre-miRNAs (K1 to K12). (D) Expression of XPO1 following siRNA knockdown analyzed by Western-blotting in MM and KMM cells. (E) Expression of XPO1 following siRNA knockdown analyzed by Western-blotting in AGS and HUH7 cells. (F) Analysis of cell proliferation following siRNA knockdown of XPO1 in MM, KMM, AGS and HUH7 cell lines. (G) Analysis of cell proliferation following treatment with DMSO or KPT-8602 at different concentrations for 3 days in MM, KMM, AGS and HUH7 cells. (H) Formation of colonies in soft agar following treatment with DMSO or KPT-8602 at 1 µM in KMM, AGS and HUH7 cells. Representative fields are shown and efficiencies of colony formation are presented.

We performed RNA interference knockdown to confirm the results obtained in CRISPR-Cas9 screening in KMM cells as well as cell lines of other types of cancer including AGS and HUH7 derived from gastric and liver cancers, respectively. The role of XPO1 in gastric cancer has not been examined before and has only been minimally examined in liver cancer(25). Knockdown of XPO1 decreased cell proliferation in KMM, AGS and HUH7 cells but had no effect on the primary MM cells (Fig. 3D-F). Using a XPO1 specific SINE compound KPT-8602, we observed a significant inhibition of cell proliferation at a dose-dependent fashion starting at 0.1 µM in KMM, and 0.5 µM in AGS and HUH7 cells (Fig. 3G). MM cells were only sensitive to KPT-8602 at concentrations >1 µM. Furthermore, inhibition of XPO1 abrogated colony formation of KMM, AGS and HUH7 cells in softagar (Fig. 3H), indicating the essential role of XPO1 in maintaining cellular transformation.

### Inhibition of XPO1 induces p53 activation and cell cycle arrest

We examined the molecular mechanism mediating the inhibition of cell proliferation by KPT-8602. Treatment of MM, KMM, AGS, and HUH7 cells with KPT-8602 at 1 µM for 24 h induced cell cycle arrest at G2/M phase in MM and KMM, and at G0/G1 phase in AGS and HUH7 cells (Fig. 4A), suggesting different responses of these cells to KPT-8602. In parallel, we observed an activation of the p53 pathway as shown by the increase of phospho-p53 in KMM, AGS and HUH7 cells but not MM cells after treatment with 1 µM of KPT-8602 for 24 h (Fig. 4B). p53 activation has been shown to be dependent on PML-NBs within the nucleus. Interestingly, both MM and KMM cells had strong nuclear PML staining as well as cytoplasmic staining with some cells showing staining pattern similar to PML-NBs, and KPT-8602 treatment did not alter the PML staining pattern in these cells (Fig. 4C). These results indicated that PML-NBs were unlikely to mediate p53 activation in KMM cells. In contrast, AGS cells had no nuclear PML staining and only some HUH7 cells had weak nuclear PML staining, none of which showed staining pattern of PML-NBs (Fig. 4C). However, strong nuclear staining with PML-NBs pattern was observed following KPT-8602 treatment in AGS and HUH7 cells (Fig. 4C). Significantly, siRNA knockdown of PML (Fig. 4D), which disrupted PML-NBs, inhibited KPT-8602 induced p53 phosphorylation in AGS and HUH7 cells, highlighting the essential role of PML-NBs in p53 activation in these cells (Fig. 4E).

**FIG 4.**
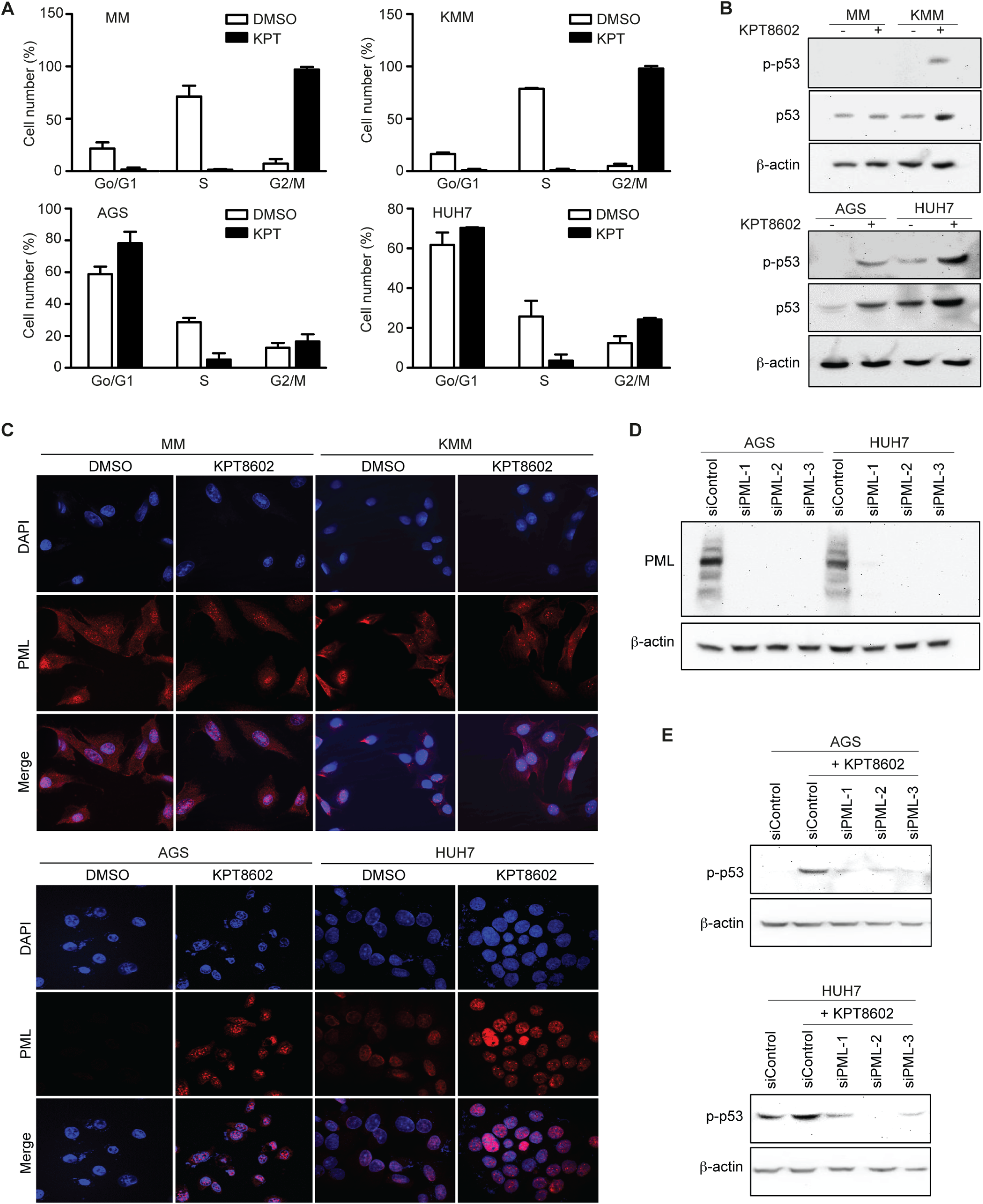
Inhibition of XPO1 induces PML-mediated p53 activation and cell cycle arrest. (A) Effect of KPT-8602 treatment on cell cycle progression of MM, KMM, AGS and HUH7 cells. Cells treated with 1 µM of KPT-8602 for 48 h were analyzed by flow cytometry after BrdU and propidium iodide staining. (B) Analysis of phospho-p53 (p-p53) and total p53 in MM, KMM, AGS and HUH7 cells after treatment with 1 µM KPT-8602 by Western-blotting. (C) Examination of PML in MM, KMM, AGS and HUH7 cells after treatment with 1 µM KPT-8602 for 24 h by immunofluorescence assay. The slides were counterstained with DAPI, and pictures were taken with a confocal microscopy (magnification x600). (D) Expression of PML in AGS and HUH7 cells following siRNA knockdown analyzed by Western-blotting. (E) Analysis of p-p53 in AGS and HUH7 cells following siRNA knockdown of PML analyzed by Western-blotting.

### p53 activation in PML-NBs depends on the nuclear accumulation of autophagy adaptor protein p62 (SQSTM1) in AGS and HUH7 cells

Since PML-NBs are thought to form hybrid bodies with p62 protein during cellular stress(26), we investigated the p62 protein after XPO1 inhibition in AGS and HUH7 cells. Treatment with 1 µM KPT-8602 for 24 h increased the expression level of p62 in AGS and HUH7 cells (Fig. 5A). Interestingly, we observed nuclear translocation and accumulation of p62, which was colocalized with p53 in AGS and HUH7 cells after KPT-8602 treatment (Fig. 5B). Furthermore, p62 was colocalized with PML-NBs in AGS and HUH7 cells following XPO1 inhibition, which was consistent with the hypothesis of formation of hybrid bodies (Fig. 6A). The expression level of p62 was also increased in KMM cells but not in MM cells (Fig. 5A). However, we did not observe nuclear translocation and accumulation of p62 in MM and KMM cells, and there was no nuclear colocalization of p53 with p62 in these cells (Fig. 5B). Consistent with these results, there was no obvious nuclear colocalization of p62 with PML-NBs in MM and KMM cells (Fig. 6A). These results indicated that p62 was unlikely to be involved in p53 activation following XPO1 inhibition in KMM cells.

**FIG 5.**
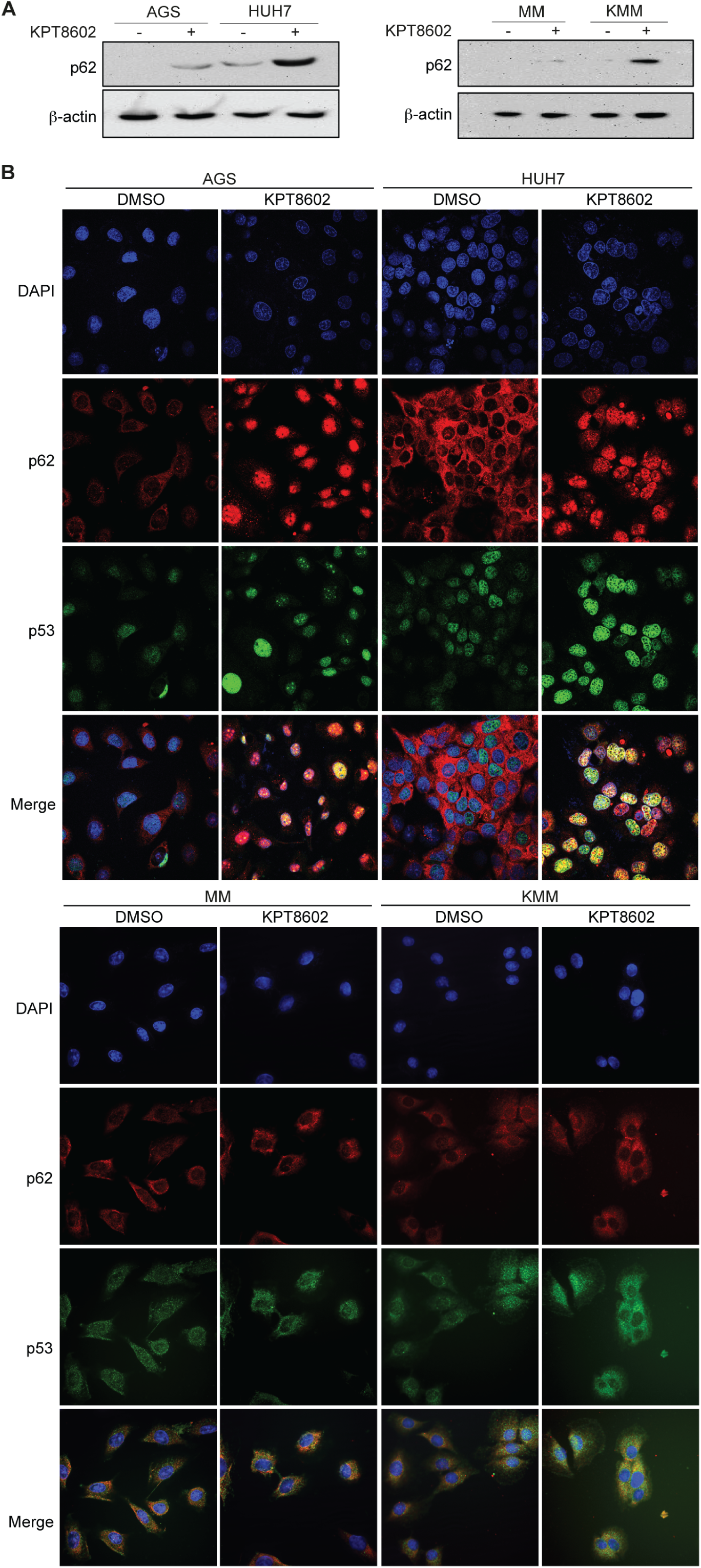
Induction of p62 nuclear accumulation and colocalization with p53 following XPO1 inhibition in AGS and HUH7 cells. (A) Expression of p62 in MM, KMM, AGS and HUH7 cells after treatment with 1 µM KPT-8602 for 24 h analyzed by Western-blotting. (B) Expression of p53 and p62 in AGS, HUH7, MM and KMM cells after treatment with 1 µM KPT-8602 for 24 h analyzed by immunofluorescence assay. The sections were counterstained with DAPI, and pictures were taken with a confocal microscopy (magnification x600).

**FIG 6.**
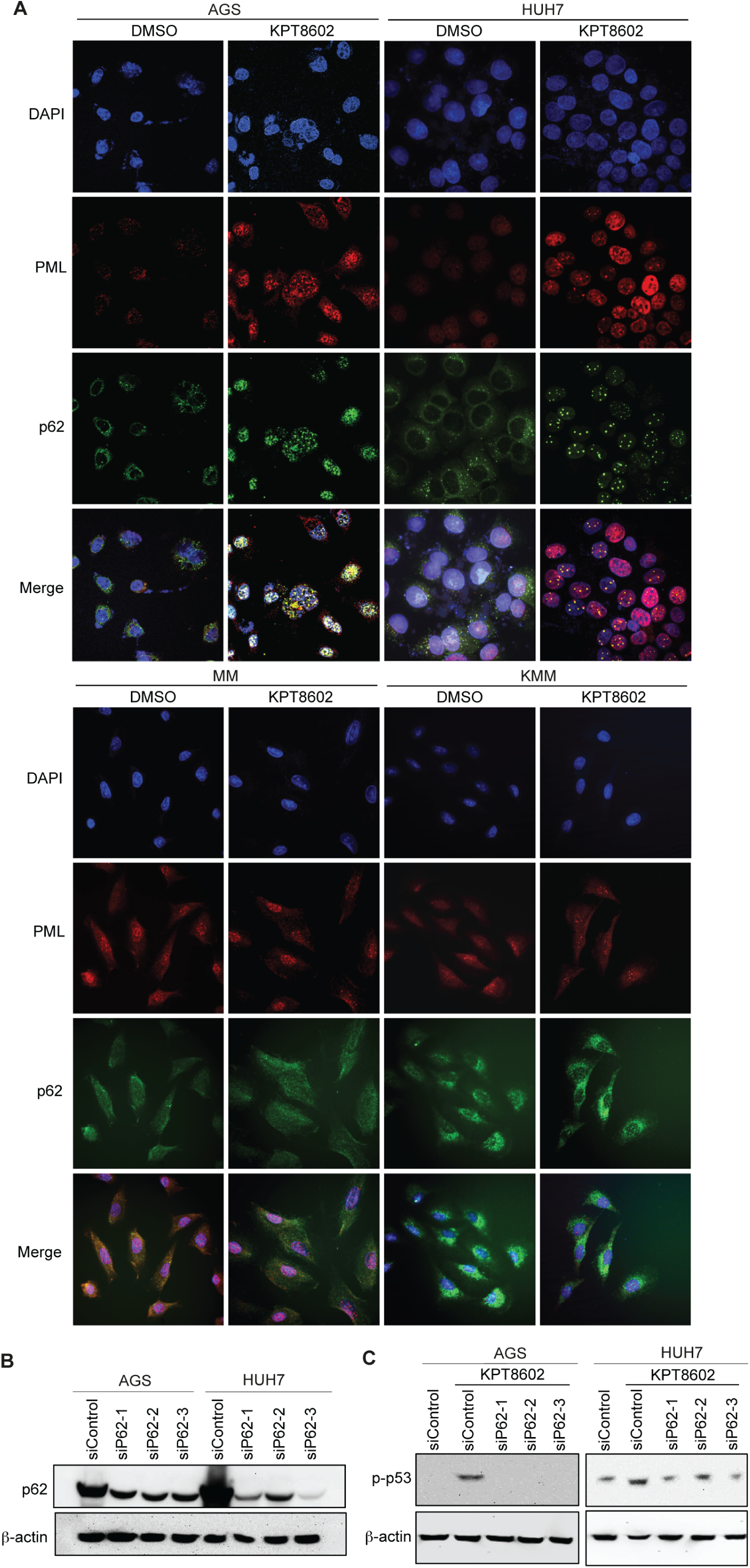
XPO1 inhibition-induced p53 activation in PML-NBs depends on p62 nuclear accumulation in AGS and HUH7 cells. (A) Expression of PML and p62 in AGS, HUH7, MM and KMM cells after treatment with 1 µM KPT-8602 for 24 h analyzed by immunofluorescence assay. The sections were counterstained with DAPI, and pictures were taken with a confocal microscopy (magnification x600). (B) Expression of p62 in AGS and HUH7 cells following siRNA knockdown analyzed by Western-blotting. (C) Analysis of p-p53 in AGS and HUH7 cells after treatment with 1 µM KPT-8602 for 24 h following siRNA knockdown of p62 by Western-blotting.

To investigate the role of p62 in p53 activation in AGS and HUH7 cells, we performed knockdown of p62 in AGS and HUH7 cells (Fig. 6B). Knockdown of p62 abolished p53 activation in AGS cells and significantly decreased p53 activation in HUH7 cells following treatment with KPT-8602 (Fig. 6C). These results indicate an essential role of p62 accumulation in XPO1 inhibition-induced p53 activation in PML-NBs in AGS and HUH7 cells.

## Discussion

Numerous genetic engineering techniques using endonucleases to induce double-stranded breaks (DSBs) at specific sites in the target DNA have been developed in the last few decades. Endonucleases, such as transcription activator-like effector nucleases (TALENs) or zinc finger nucleases (ZFNs), were previously used to inactivate targeted genes but some studies highlighted their lack of specificity and poor efficiency(27). More recently, CRISPR-Cas9 has been demonstrated as an efficient and precise tool for genetic disruption or correction in vitro and in vivo(10, 11).

In this report, by performing a genome-wide CRISPR-Cas9 screening, we characterized gene essentiality in KSHV-transformed cells(13). KSHV, as a human oncogenic virus in the family of Herpesviridae, is associated with several human malignancies, including KS, primary effusion lymphoma (PEL), KSHV inflammatory cytokine syndrome (KICS) and a subset of multicentric Castleman’s disease (MCD)(28). Using a model of KSHV-induced cellular transformation of primary cells(13), we have identified essential genes in both MM and KMM cells, and observed an increase in survival ability in KSHV-transformed cells compared to primary cells in a genome-wide CRISPR-Cas9 knockout screening (Fig. 1), confirming the oncogenic nature of the virus. Moreover, we have identified genes involved in the survival of KMM or MM cells (Fig. 2). In parallel, we have observed a significant positive correlation in gene essentiality between KMM cells and other cancer cells observed by similar CRISPR-Cas9 screening(17), illustrating the convergence of common survival features in these cancer cells.

Current therapies targeting latent infections of oncogenic viruses are often limited in efficacy, and cannot eradicate the viruses. Numerous studies have performed CRISPR-Cas9 screening in viral cancer models to identify novel therapeutic targets(29). CRISPR-Cas9 genome editing in models of oncogenic viruses should facilitate the identifications of vulnerable genes/mutations, viral and cellular oncoproteins, and essential or restriction factors for infections. Particularly, in the case of KSHV, CRISPR-Cas9-mediated gene knockout enabled the identification of RSK, an important substrate of viral lytic protein ORF45 required for KSHV gene expression and production of virions(30, 31). Recently, a CRISPR-Cas9 screening in 8 PEL cell lines has been performed(32). They identified 210 essential genes across these cell lines and highlighted the dependence of IRF4 and MDM2 pathways in PEL survival. They also showed the dependency of PEL cells on cyclin D2 and c-Flip. In our study, we have used matched primary MM cells as controls and identified 1,259 genes (Group 8) that are essential for maintaining the proliferation of KMM cells, of which, 18 genes had CRISPR score ratios of <-5 (−32 fold) at day 21 over day 1 for KMM cells (Fig. 2). Among the top enriched pathways are those that are related to the mTOR pathway, which has been shown to be the most effective target in KS patients in clinical studies(18), thus validating the relevance of the model. These enriched pathways are likely vulnerable targets of KSHV-induced malignancies. In agreement with the PEL screening, we have identified IRF signaling as one of the Top 10 enriched pathways with genes losing sgRNAs in KMM over MM cells; however, the hypoxia signaling pathway and p38 MAPK pathway are the top enriched pathways with genes losing sgRNAs. These pathways have putative suppressive effects, and hence are likely essential for maintaining the homeostasis of KSHV-transformed cells.

The current study has identified XPO1 as a critical factor for the proliferation of KSHV-transformed cells. We confirmed the essential role of XPO1 in cell proliferation and cellular transformation of cell lines derived from gastric and liver cancers (Fig. 3). XPO1 dysregulation affects fundamental cellular processes such as inflammatory responses, cell cycle and apoptosis, and might contribute to tumorigenesis(2). Since numerous tumor suppressor proteins and oncoproteins, including p53, APC, Rb, NFAT, FOXOs, p27, nucleophosmin, BCR-ABL, eIF4E, survivin and β-catenin, harbor NES, and are exported from the nucleus to the cytoplasm by XPO1 to fulfill their growth-promoting and anti-apoptotic functions, nuclear-cytoplasmic transport, particularly those mediated by XPO1, is likely an effective general target for cancer(2).

Upregulation of XPO1 is observed in many types of cancer tissues such as lung cancer, osteosarcoma, pancreatic cancer, ovarian cancer, cervical carcinoma, gastric and hepatocellular carcinoma, myeloid and lymphoid leukemia, mantle cell lymphoma, and multiple myeloma(22). In parallel, XPO1 upregulation has been shown to affect the functions of several oncogenes such as VEGF, EGF receptor, Cox-2, c-Myc and HIF-1, which are not the direct cargos of XPO1(33). In addition to XPO1 upregulation, post-translational modifications of oncogenes and tumor suppressors can also alter or enhance their nuclear-cytoplasmic transport by XPO1. For example, p53 sumoylation and ubiquitination have been shown to enhance its XPO1-mediated export(34, 35). Drug resistance and poor prognosis have also been correlated with XPO1 upregulation in several malignancies(36, 37). For example, XPO1 upregulation induces aberrant nuclear-cytoplasmic export of Topoisomerase IIα (Topo IIα) involved in DNA replication, transcription, and chromatin segregation. Topo IIα is a target of numerous anticancer drugs(38). However, its cytoplasmic export often induces resistance to Topo IIα-specific inhibitors such as doxorubicin and etoposide. As a result, XPO1 inhibition increases sensitivity to Topo IIα inhibitor(38). Hence, dual treatment with both Topo IIα inhibitor and XPO1 inhibitor is likely to improve the efficacy of cancer therapy.

The mechanism of XPO1 upregulation in cancer cells is still not well understood. Chromosomal translocations, and mutations or gain of copies of XPO1 gene could explain XPO1 overexpression. Indeed, a recurrent mutation in codon 571 of XPO1 gene has been reported in acute lymphoblastic leukemia (ALL) and in an adult acute myeloid leukemia (AML) with *CBL* syndrome displaying XPO1 overexpression(39, 40). Moreover, Myc and p53, often found modulated in cancer cells, have been shown to positively and negatively alter the expression of XPO1, respectively(41). Finally, a copy number gain in the XPO1 locus was also observed in patients affected by primary mediastinal B-cell lymphoma (PMBL) and diffuse large B-cell lymphoma (DLBCL)(42, 43). We observed increased XPO1 expression in KMM cells compared to MM cells, and that both vCyclin and pre-miR-K3 mediated XPO1 upregulation (Fig. 3A-3C). It would be interesting to investigate the effect of cell cycle progression on XPO1 upregulation.

SINE compounds have been used to treat several types of cancer, among them, KPT-330 is currently in Phase 1 and 2 clinical trials in numerous solid and hematologic malignancies with and without in combination with other chemotherapeutic agents, and has shown promising results(33). To decrease adverse side effects, a second generation of SINE compounds including KPT-8602 with improved tolerability have been developed and tested in clinical trials(44). In this report, we have shown that KPT-8602 inhibits the cell proliferation and cellular transformation of KMM, AGS and HUH7 cells (Fig. 3), confirming the anti-cancer effect of SINE family. Indeed, KPT-8602 has been shown to inhibit the proliferation of leukemia cells derived from ALL in vitro and increase the animal survival rate in ALL xenograft models(45).

SINE compounds can block XPO1-mediated export, and therefore cause mislocalization of tumor suppressors or oncoproteins, and decrease the survival of cancer cells(1). A previous study highlighted the nuclear accumulation of p21, p27 and FOXO proteins after treatment with SINE compound S109 in colorectal cancer cells(46). Recently, KPT-330 has been shown to induce p53 and p21 retention within the nucleus in gastric cancer cells(47). In our report, we have observed cell cycle arrest associated with p53 activation in KMM, AGS and HUH7 cells after KPT-8602 treatment. However, we have found that different types of cancer cells respond differently to this inhibitor with KMM cells manifesting G2/M arrest, and AGS and HUH7 cells manifesting G0/G1 arrest (Fig. 4A). Consistent with these results, the mechanism mediating p53 activation in AGS and HUH7 cells is different from that in KMM cells. We have observed p53 nuclear accumulation in AGS and HUH7 cells but not KMM cells after KPT-8602 treatment (Fig. 5B).

PML-NBs recruit a variety of proteins and modulate their posttranslational modifications. Particularly, p53 and its regulators (ARF, HIPK2, CBP, MDM2, SIRT1 and MOZ) traffic through PML-NBs, suggesting that these nuclear bodies could regulate p53 activation by posttranslational modifications(48). Indeed, disruption of PML-NBs inhibits p53 activation and induces expression of its downstream genes, such as Bax and p21(9, 49, 50); however, the underlying mechanism remains unclear. Interestingly, we have confirmed that p53 nuclear accumulation depends on the formation of PML-NBs after XPO1 inhibition in AGS and HUH7 cells (Fig. 4C-E). On the other hand, p53 activation in KMM after XPO1 inhibition does not depend on the formation of PML-NBs, which remains to be further explored.

Other NBs have been identified in the nucleus, and some studies suggested possible fusion events among them, forming hybrid NBs(51). Interestingly, XPO1 is involved in the formation of CRM1-nucleolar bodies (CNoBs), and its inhibition by XPO1 inhibitor Leptomycin B disrupts CNoBs(52). Another type of NBs involved with p62 (SQSTM1) was identified by imaging(26). p62 protein is mostly described as a scaffold cytoplasmic protein involved in autophagy processes by interacting with LC3, and therefore mediating the recruitment of LC3 to poly-ubiquitinated protein aggregates to form autophagosome in the cytoplasm(53). Unlike its cytoplasmic function, the role of p62 within the nucleus is still not well understood. While PML-NBs are ubiquitous, CNoBs and p62-NBs are stress-induced. It has been reported that p62-NBs are colocalized with PML-NBs upon inhibition of CNoBs(51), highlighting the role of nuclear p62 in the recruitment of poly-ubiquitinated proteins to PML-NBs(26).

The role of p62 in cancer development is still controversial(54). The expression level of p62 can be increased in cancer cells, and its overexpression has been shown to enhance cellular transformation through the activation of NRF2, mTORC1 and c-Myc pathways in liver cancer cells, independent of the autophagy pathway(55). On the other hand, in a non-tumorigenic environment, p62 attenuates inflammation and fibrosis(56), and p62 loss has been shown to increase tumorigenesis in epithelial cells(57).

In this study, we have demonstrated the essential role of p62 in the activation of p53 tumor suppressor after XPO1 inhibition in AGS and HUH7 cells (Fig. 5 and 6). We have observed nuclear accumulation of p62, which is colocalized with p53 after XPO1 inhibition in AGS and HUH7 cells. In parallel, we have observed colocalization of p62 with PML-NBs, hence enforcing the hypothesis of formation of PML-p62 hybrid NBs in stress conditions(26). Finally, we have demonstrated for the first time that nuclear accumulation of p62 is required for p53 phosphorylation and activation within the PML-NBs after KPT-8602 treatment in in AGS and HUH7 cells, demonstrating the indirect role of p62 in regulating the function of p53 in these cancer cells. Interestingly, while XPO1 inhibition causes an increase of p62 level in KMM cells, it neither leads to p62 nuclear accumulation nor mediates p53 activation (Fig. 5 and 6). It would be interesting to investigate whether the increase of p62 level after XPO1 inhibition contributes to the G2/M cell cycle arrest in KMM cells.

## Materials and Methods

### Cells and inhibitor

Rat primary embryonic mesenchymal stem cells (MM), KSHV-transformed MM cells (KMM), AGS and HUH7 cell lines were maintained in Dulbecco modified Eagle medium (DMEM) supplemented with 10% fetal bovine serum (FBS; Sigma-Aldrich), 4 mM L-glutamine, and 10 µg/ml penicillin and streptomycin. KPT-8602 was obtained from Selleckchem.

### Rat sgRNA library cloning

Rat sgRNA library cloning was carried out as previously described(58). This sgRNA library consists in a library of 59,520 unique sgRNAs targeting 19,840 coding genes, each with 3 independent sgRNAs, and 10 non-targeting sgRNAs, for a total of 59,530 sgRNAs. Specifically, we carefully chose three target sites for each gene to cover most common exon in transcript variants ranging from 3% to 70% in coding sequence (CDS) region from the rat genome (Rattus novegicus; Rnor 5.0 version). Furthermore, we chose the target sites by avoiding potential off-target effects and in-frame mutations using Cas-OFFinder(59) and Cas-Designer(60) (http://www.rgenome.net/).

### Lentiviral production of the sgRNA library

Lentiviral production was carried out as previously described(61, 62). Briefly, HEK293T cells, were seeded at ∼40% confluence one day before transfection in DMEM supplemented with 10% serum. One-hour prior to transfection, DMEM media was removed and fresh OptiMEM media (Life Technologies) was added. Lipofectamine 2000 (Life Technologies) was used to transfect 20 µg of lentiCRISPR plasmid library, 10 µg of pVSVg and 15 µg of psPAX2. Lipofectamine 2000 of 100 µl was diluted in 4 ml of OptiMEM, and after 5 min, it was added to the mixture of plasmid DNAs, incubated for 20 min and then added to the cells. Media was refreshed after 6 h and collected after 3 days. The supernatant was centrifuged at 3,000 rpm at 4°C for 10 min to pellet cell debris, filtered (0.45 µm), and concentrated by ultracentrifugation (Beckmann) at 24,000 rpm for 2 h at 4°C. The virus preparation was finally resuspended overnight at 4°C in DMEM medium, aliquoted and stored at −80°C.

### Lentiviral transduction of the sgRNA library

Lentiviral transduction was carried out as previously described(61). Cells were transduced with the shRNA library via spinfection at a MOI of 0.3 in full DMEM medium supplemented with 10% fetal bovine serum, 4 mM L-glutamine, and 10 µg/ml penicillin and streptomycin in the presence of 10 µg/ml of polybrene. Flasks containing cells were centrifuged at 2,000 rpm for 2 h at 37°C. After the spin, media was removed and fresh DMEM media was added to the cells.

### Genome DNA sequencing

Genome DNA sequencing was carried out as previously described(10). Genomic DNA was extracted using QiAMP kit (Qiagen) and PCR was performed in two steps. First, the input genomic DNA was amplified in order to achieve 300X coverage over the sgRNA library using the primers F1: TCG TCG GCA GCG TCA GAT GTG TAT AAG AGA CAG TAT CTT GTG GAA AGG ACG AAA and R1: GTC TCG TGG GCT CGG AGA TGT GTA TAA GAG ACA GTT ATT TTA ACT TGC TAT TTC TAG CTC. Then, to attach Illumina adaptors and barcode samples, a second PCR reaction was carried out using the primers F2: AAT GAT ACG GCG ACC ACC GAG ATC TAC ACT CTT TCC CTA CAC GAC GCT CTT CCG ATC T, and R2: CAA GCA GAA GAC GGC ATA CGA GAT GTG ACT GGA GTT CAG ACG TGT GCT CTT CCG ATC T. Amplicons from the second PCR were gel-extracted, quantified, mixed and sequenced using HiSeq 2500 (Illumina).

### Ingenuity pathway analysis

The differentially enriched pathways at day 21 over day 1 post-transduction in KMM over MM cells were evaluated using IPA software (Ingenuity H Systems, USA; http://www.ingenuity.com).

### Soft agar assay

Soft agar assay was carried out as previously described(15).

### Cell proliferation assay

MM, KMM, AGS and HUH7 cells plated at a density of 200,000 cells/well and treated with DMSO or KPT-8602 at different concentrations were counted daily using a Malassez chamber.

### Western blot

Western-blotting was carried out as previously described(62). Primary antibodies to β-actin (Santa Cruz), p62 (CST), SIRT1 (CST), Cas9-HA (Santa Cruz), PML (CST), XPO1 (CST), p53 (CST), and phospho-p53 (CST) were used.

### Immunofluorescence assay

Cells were fixed in methanol for 10 minutes at room temperature and processed for antibody staining as previously described(63). Immunostaining was performed using anti-p62 antibody (CST), anti-PML antibody (CST) or anti-phospho-p53 antibody (CST). Alexa488-and Alexa568-conjugated secondary antibodies (Thermo Fisher Scientific) were used to reveal the signals. Nuclei were counterstained with DAPI. Tissue sections without incubation with primary antibodies were used as negative controls. Images of representative areas were acquired using a confocal fluorescence microscopy with a 60x objective (Nikon Eclipse C1).

### Cell cycle assay

The cell cycle was analyzed as previously described(62). MM, KMM, AGS and HUH7 cells pulsed with 10 µM 5-bromo-2′-deoxyuridine (BrdU) (B5002; Sigma-Aldrich) were stained with propidium iodide (P4864; Sigma-Aldrich). BrdU was detected by flow cytometry with a Pacific Blue-conjugated anti-BrdU antibody (B35129; Thermo Fisher Scientific). Results were analyzed using FlowJo software (FlowJo LLC, USA).

### Statistical analysis

Statistical analysis was performed using the Kolmogorov-Smirnov test or the two-tailed t test as indicated in the figure legends, and a P value of ≤0.05 was considered significant. Single, double, and triple asterisks in figures represent P values of ≤0.05, ≤0.01, and ≤0.001, respectively, while NS indicates not significant.

### Data assess

All CRISPR data generated in this study have been submitted to the NCBI Gene Expression Omnibus and will become publically available with accession number GSE125507.

## Acknowledgments

This work was supported by grants from the National Institute of Health (CA096512, CA124332, CA132637, CA177377, CA213275, DE025465 and CA197153 to S-J Gao). National Natural Science Foundation of China (81730062 and 81761128003 to C Lu) and Nanjing Medical University (KY101RC1710 to C. Lu). We thank members of Dr. Gao’s laboratory for technical assistances and helpful discussions.

## Supplemental Figure Legends

**FIG S1.** (A) Expression of Cas9 protein in MM and KMM cells stably expressing Cas9 analyzed by Western-blotting. (B) Expression of SIRT1 protein following CRISPR-Cas9 knockout analyzed by Western-blotting in Cas9-expressing MM and KMM cells. (C) Formation of colonies in soft agar of Cas9-expressing MM and KMM cells. Representative fields are shown.

**FIG S2.** Correlation between mRNA expression determined by RNA-seq(16) and gene essentiality based on CRISPR score in MM and KMM cells.

**FIG S3.** Analysis of relative sgRNA counts at days 1, 4, 11, and 21 post-transductions for XPO1, XPO2, XPO3, XPO4, XPO5, XPO6 and XPO7 genes in MM and KMM cells.

## Supplemental Tables

**Table S1.** Different groups of genes identified by Crispr-Cas9 screening of MM and KMM cells.

**Table S2.** List of depleted pathways in KMM over MM cells at day 21 versus day 1 post-transduction analyzed by IPA.

**Table S3.** List of enriched pathways in KMM over MM cells at day 21 versus day 1 post-transduction analyzed by IPA.

